# Growth Patterns of birds, dinosaurs and reptiles: Are differences real or apparent?

**DOI:** 10.1101/597260

**Authors:** Norbert Brunner, Manfred Kühleitner, Werner-Georg Nowak, Katharina Renner-Martin, Klaus Scheicher

**Affiliations:** University of Natural Resources and Life Sciences, Department of Integrative Biology and Biodiversity Research, A-1180 Vienna, Austria

**Author notes:** **Corresponding author:** Manfred Kühleitner, Institute of Mathematics, Department of Integrative Biology and Biodiversity Research, University of Natural Resources and Life Sciences (BOKU), Gregor Mendel Strasse 33, A-1180 Vienna, Austria, **E-Mail:**.

**Keywords:** Bertalanffy-Pütter differential equation, *Tenontosaurustilletti*, *Alligator mississippiensis*, Athens Canadian Random Bred strain of *Gallus gallusdomesticus*

## Abstract

Systematics of animals was done on their appearance or genetics. One can also ask about similarities or differences in the growth pattern. Quantitative studies of the growth of dinosaurs have made possible comparisons with modern animals, such as the discovery that dinosaurs grew in relation to their size faster than modern reptiles. However, these studies relied on only a few growth models. If these models are false, what about the conclusions? This paper fits growth data to a more comprehensive class of models, defined by the von Bertalanffy-Pütter differential equation. Applied to data about dinosaurs, reptiles and birds, the best fitting models confirmed that dinosaurs may have grown faster than alligators. However, compared to modern broiler chicken, this difference was small.

## 1. INTRODUCTION

Mathematical growth models aim at a simplified description of growth in terms of curves that fit well to size-at-age data [1]. As the growth of animals depends on multiple factors, the most-informative data came from controlled studies, where animals were reared under the same conditions and weighed repeatedly during the entire phase of growth. This was feasible e.g. for chicken [2]. By contrast, for wildlife and wild-caught fish, in general for each animal there was only one measurement of mass-at-age. Even with data about thousands of animals there remained considerable uncertainties about the proper choice of the growth model [3]. For extinct species the situation was even worse, as no weighing of body mass was possible for fossils. However, recent approaches led to mathematical growth models for dinosaurs [4] and thereby to a comparison of growth pattern of different species. These quantitative studies have “revolutionized our understanding of dinosaur biology” [5].

Growth studies for vertebrates relied on few models only. Examples are the models of Brody [6], von Bertalanffy [7], Gompertz [8], Richards [9, 10], West [11], Verhulst [12] logistic growth, and the generalized Bertalanffy model promoted by Pauly [13].This paper studies the comprehensive class of growth models (1).

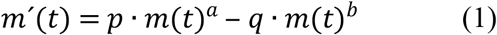

It describes growth about mass *m*(*t*) at time *t* and it uses five model parameters, namely the non-negative exponent-pair *a* < *b*, the constants *p* and *q*, and the initial mass *m*(0) = *m*_0_ > 0:

Equation (1) was proposed by Pütter [14] and von Bertalanffy [15]. As shown in Figure 1, the above-mentioned named models are special cases of it, whereby each model corresponds to a different exponent-pair or to a line segment of exponent-pairs. The Gompertz model is a limit case on the diagonal. In view of the exceptional character of the named models, we ask, if there are other models from the Bertalanffy-Pütter class that describe growth pattern of dinosaurs better and thereby allow for more accurate comparisons between different species.

**Figure 1.**
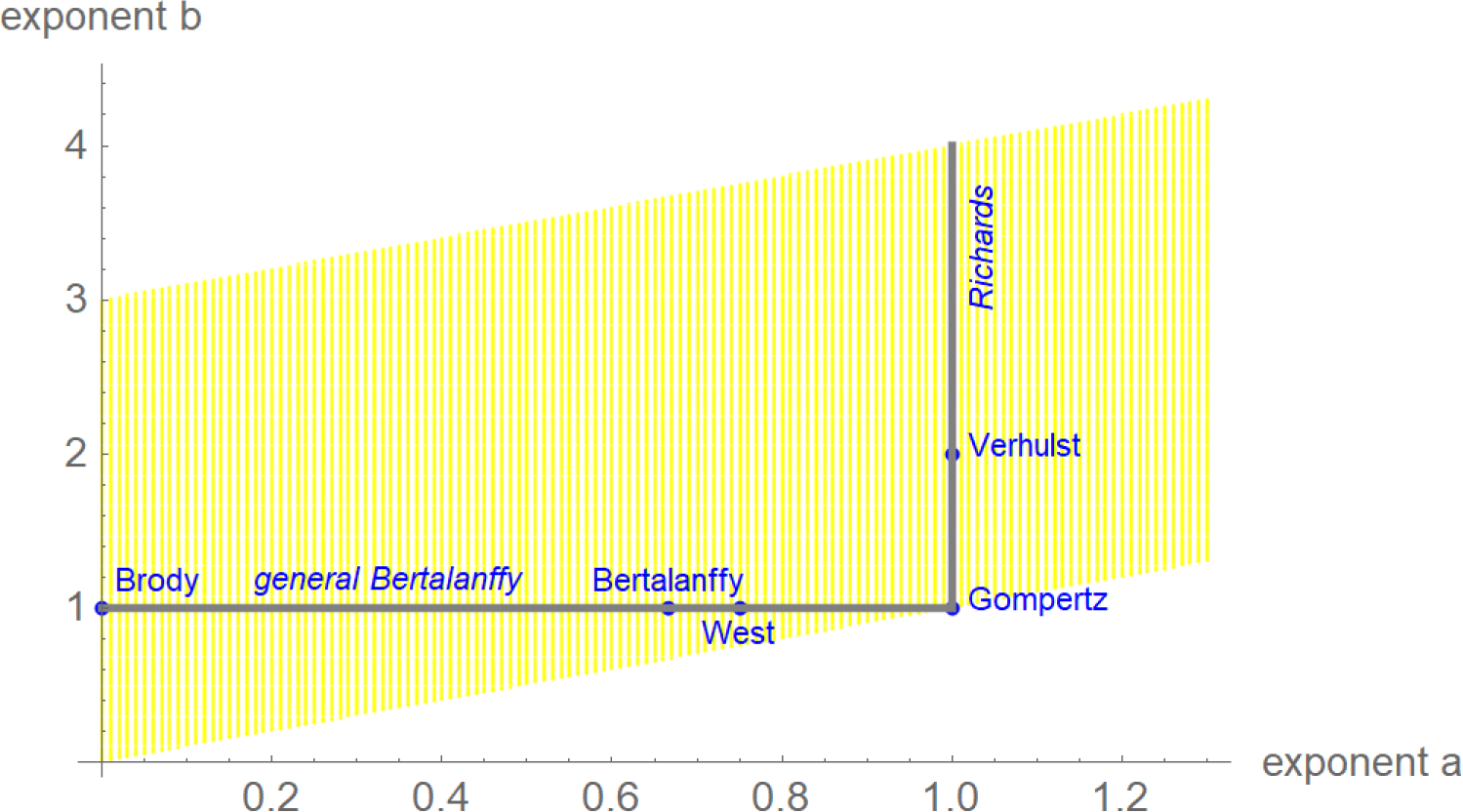
Named models (blue) and part of the search-region (yellow) for the exponent-pair of the best fitting growth model.

We illustrate these questions by a case study, where we identify growth models from the general class (1) with the best fit to mass-at-age data for a species of dinosaurs (*Tenontosaurus*) and for two modern species of reptiles (alligators) and birds (broiler chicken) that are often compared with dinosaurs. The data were drawn from literature. In view of the need to optimize five parameters, the data-fitting problem led to an optimization problem that hitherto due to numerical instability had been almost intractable, whence practitioners confined the search for best fitting models basically to the above-mentioned named models with mathematically elementary growth curves. Recently, the authors succeeded in developing an advanced optimization method, which allowed to extend the search for the best fitting model, represented by an exponent-pair, to a much larger class of models (e.g. yellow region in Figure 1). The optimization for the present paper searched ca. 30,000-70,000 exponent-pairs (i.e. different candidate models) per data-set.

Further, in order to study the variability of the exponents, the paper identified the region of near-optimal exponent-pairs. The exponent-pairs of this region could also be used to model growth without affecting the fit to the data significantly when the other parameters were optimized.

## 2. METHODS

### Data

Mass-at-age of *Tenontosaurus tilletti* (twelve data points with mass 23-1102 kg, and age 1-26 years) was from Table 2 of [16]. Mass-at-age of *Alligator mississippiensis* (41 data-points with mass 0.1-40.7 kg and age 1-42 years) was retrieved from Figure 3A of that paper. The original source was [17], who over a time-span of forty years captured and partly recaptured ca. 7000 alligators from Loisiana, USA. Mass-at-age of *Gallus gallus domesticus* (28 data points with mass 0.04-2.23 kg and age 0-170 days) came from Table1 of [2]. This table records the average mass-at-age of 217 male chicken of the Athens Canadian Random Bred strain that survived the first 170 days since hatching. They were reared under laboratory conditions and weighed regularly.

### Materials

Data from graphics were retrieved using DigitizeIt of Bormisoft^®^. All data were copied into a spreadsheet (Excel of Microsoft^®^) and processed in Mathematica 11.3 of Wolfram Research^®^. The output of optimization was exported to a spreadsheet.

### Methods

For chicken, the best fitting growth model and the near-optimal models were identified in [18]. As the paper uses the same approach for the alligator and dinosaur data, the method is only sketched.

Assuming a lognormal distribution of mass-at-age (the standard deviation of mass is approximately proportional to mass), the maximum-likelihood model-parameters were estimated. Thereby, the method of least squares was used to fit the logarithmically transformed growth function *u*(*t*) = ln(*m*(*t*)) to the logarithmic transformation of mass data. In order to identify both the best fitting and the near-optimal exponent-pairs, for each exponent-pair on a grid the other model-parameters were optimized. Thus, using the abbreviation *SSLE*= sum of squared errors between the logarithm of the growth function and the logarithmically transformed data, the following function (2) on the grid was defined:

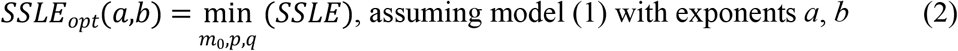

The optimization of *p*, *q*, *m*_0_ used simulated annealing, whereby for a grid point near the diagonal 50,000 annealing steps were used. For the subsequent grid points in the *b*-direction, these outputs were used as starting values and improved in 1,000 annealing steps. The output was exported to a table in the format (*a*, *b*, *m*_0_, *p*, *q*, *SSLE*_*opt*_(*a*, *b*)). It is provided as a supporting material. An exponent-pair was near-optimal, if its *SSLE*_*opt*_(*a*, *b*) exceeds the least one by less than 5%.

## 3. RESULTS

The graphical representation of the results uses red for chicken, green for alligators and blue for dinosaurs. Figure 2 plots the data and the best fitting growth curves in dimensionless coordinates. Thereby, mass is reported as a fraction of the asymptotic mass *m*_*max*_. Given the best fitting growth model, this is the limit of *m*(*t*), when time approaches infinity. Age is reported as a fraction of “full age” *t*_*full*_, at which 90% of the asymptotic mass is reached. This is used as a proxy for “adulthood”. Thereby *m*_*max*_ and *t*_*full*_ were computed from the best fitting model. Note the similarity of growth in terms of these dimensionless data.

**Figure 2.**
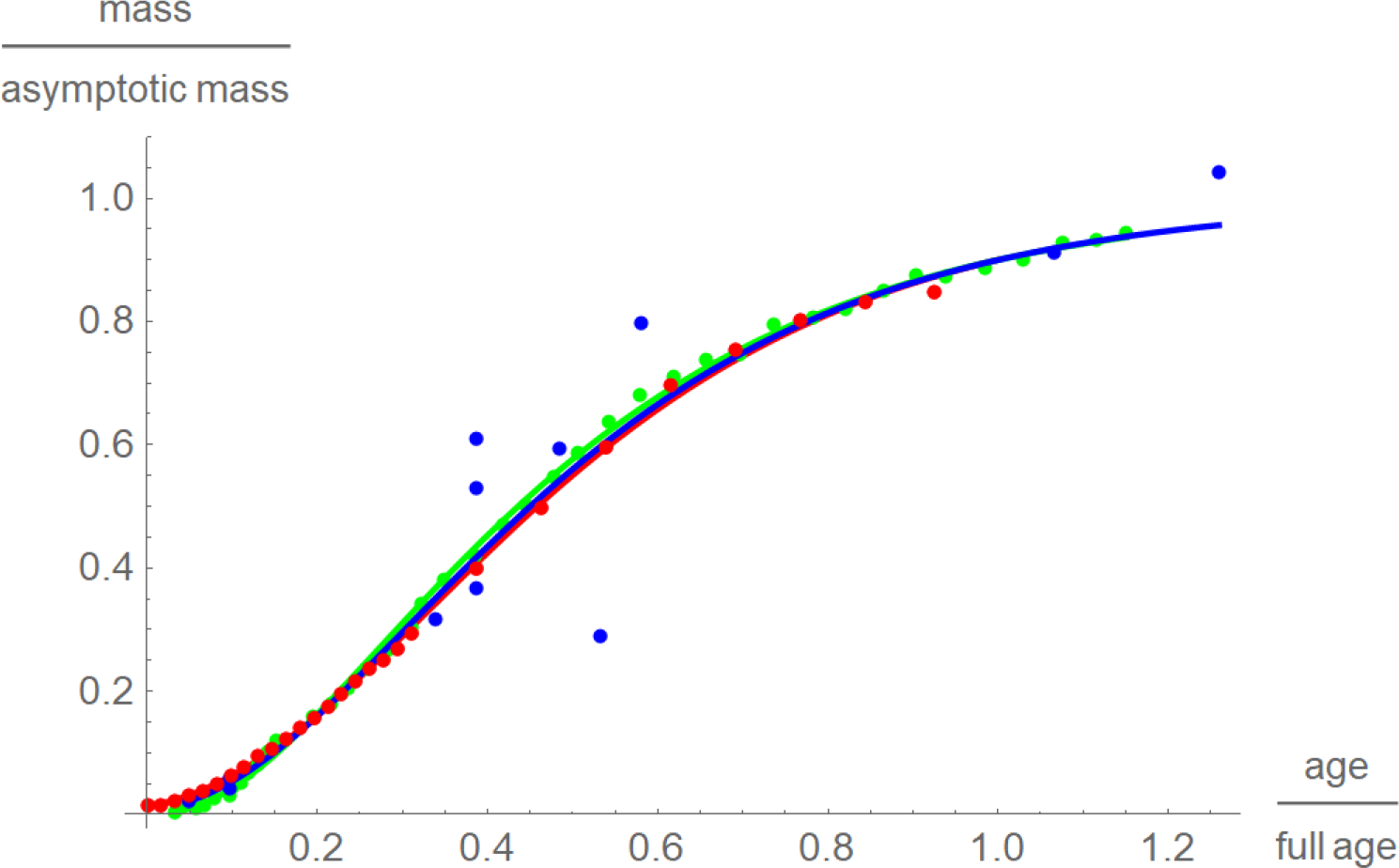
Growth data and best fitting growth curves in dimensionless coordinates (fraction of the asymptotic mass*m*_*max*_ at a fraction of the full age*t*_*full*_) for broiler chicken (red), alligators (green) and dinosaurs (blue). For chicken and alligators, but not so for dinosaurs (larger spread of the data), the data differed only slightly from the growth curves. Further, the curves were barely different.

**Figure 3.**
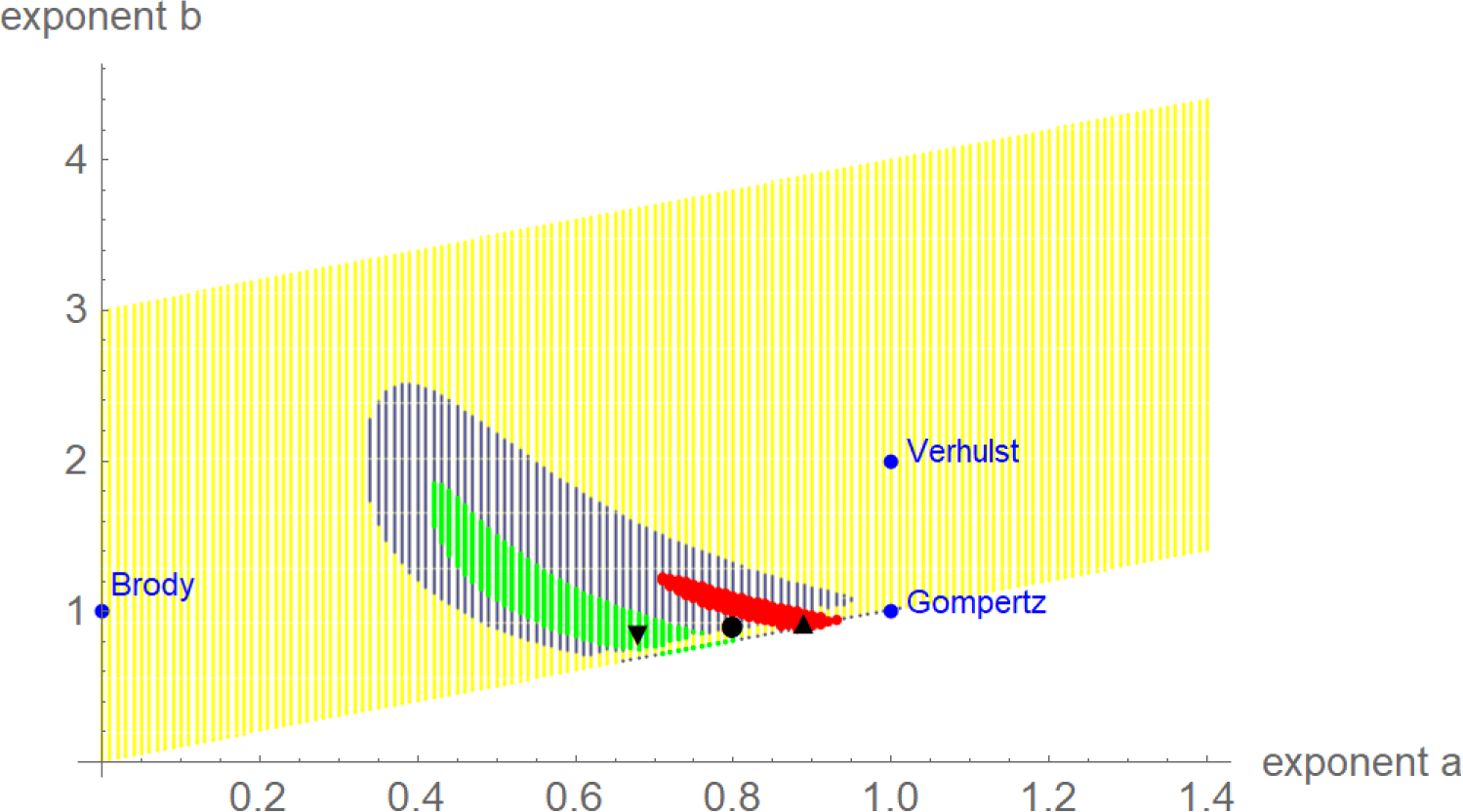
Optimal and near-optimal exponent-pairs for chicken (triangle and red area), alligators (upside triangle and green area) and dinosaurs (circle and blue area dots). For comparison with the named models, three extremal exponent-pairs are plotted (blue).

For chicken, results quoted from [18], the optimal model parameters (mass in gram, time in days) were *a* = 0.89, *b* = 0.93, *m*_0_ = 32.92 g, *p* = 1.0952, and *q* = 0.7988. This translated into an asymptotic mass of 2.67 kg, an inflection-point at day 61 with890 g (33% of the asymptotic mass) and the maximal weight gain of 19.9 g/day. For better comparison with dinosaurs, this was a maximal growth rate of 7.3 kg per year. (A dinosaur-year had more days, but these were shorter, whence overall a year covered about the same time span as today.) After 184 days (full age) 90% of the asymptotic mass was reached.

For alligators (mass in kg, time in years) the best fit was achieved for *a* = 0.68, *b* = 0.85, *m*_0_ = 158.82 g, *p* = 1.6843, and *q* = 0.8882. The asymptotic mass was 43.12 kg (slightly above the heaviest data point), the mass at the inflection point was 11.6 kg, i.e. 26% of the asymptotic mass, and the maximal growth rate was 1.78 kg/year at age 9.85 years. The full age of alligators was 36 years.

For the dinosaur-data (mass in kg, time in years) the best fit parameters were *a* = 0.8, *b* = 0.9, *m*_0_ = 22.18 kg, *p* = 6.3743, and *q* = 3.1769. The asymptotic mass was 1057.5 kg; this was slightly below the maximum mass-estimate of the data. The mass at the inflection point, 325.7 kg, was 31% of the asymptotic mass. There, at age 6.37 years, the maximal growth rate was 72.5 kg/year. Further, 90% of the asymptotic mass was reached with 21 years.

Figure 3 plots the optimal and near-optimal exponent-pairs. Despite the similarity of the data in dimensionless coordinates, the optimal exponent-pairs were different. However, due to the larger variance of the dinosaur-data the region of near-optimal exponents for dinosaurs was larger and it included both regions for alligators and chicken. Thus, judging from perspective of dinosaurs, their growth data did not display a systematic difference to modern species, whence there was no fundamental change in the growth pattern.

While these findings seem to contradict the consensus that dinosaurs grew faster than modern reptiles [5], Figure 4 compares the growth rates relative to body mass. This displays differences between the species: Well-fed broiler chicken grew more than ten times faster than alligators and dinosaurs. Further, except for a short initial period, dinosaurs grew somewhat faster than alligators. However, these comparisons were done for the best fitting model curves only.

**Figure 4.**
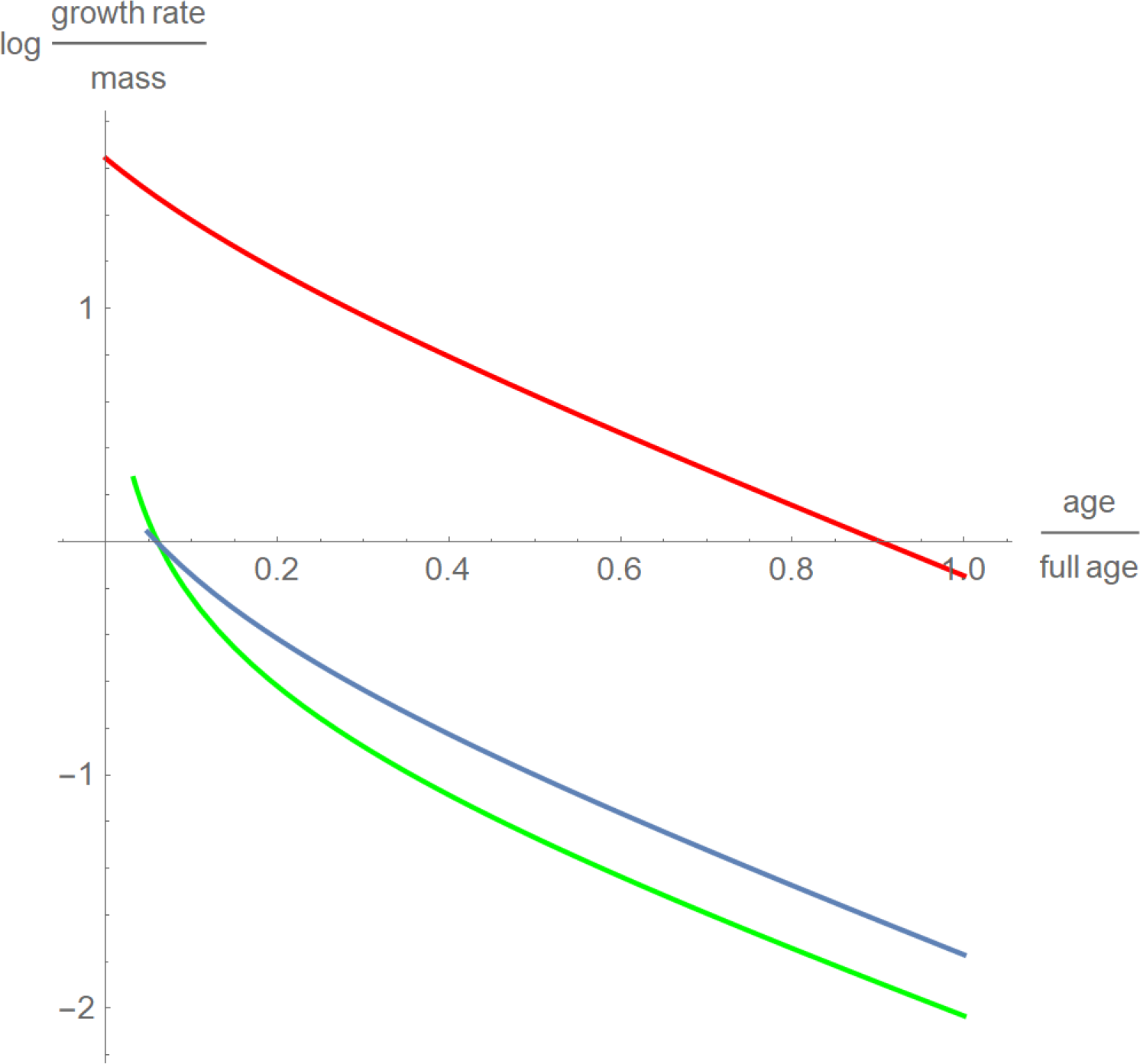
Decadic logarithm of the growth rates relative to body mass for chicken (red), alligators (green) and dinosaurs (blue) with time as a fraction of full age.

The maximal growth rate (i.e. *m’* at the inflection point) is another indicator of interest, as in comparisons between species it is used as a proxy for the basal metabolic rate [19]. Figure 5 used the near-optimal models to explore, how sensitive this indicator was to the choice of a model: The clouds were the values of *m* and *m’* at the inflection point, where *m*(*t*) was a near-optimal growth curve. Apparently, even well-fitting growth curves resulted in inaccurate estimates for the basal metabolic rate.

**Figure 5.**
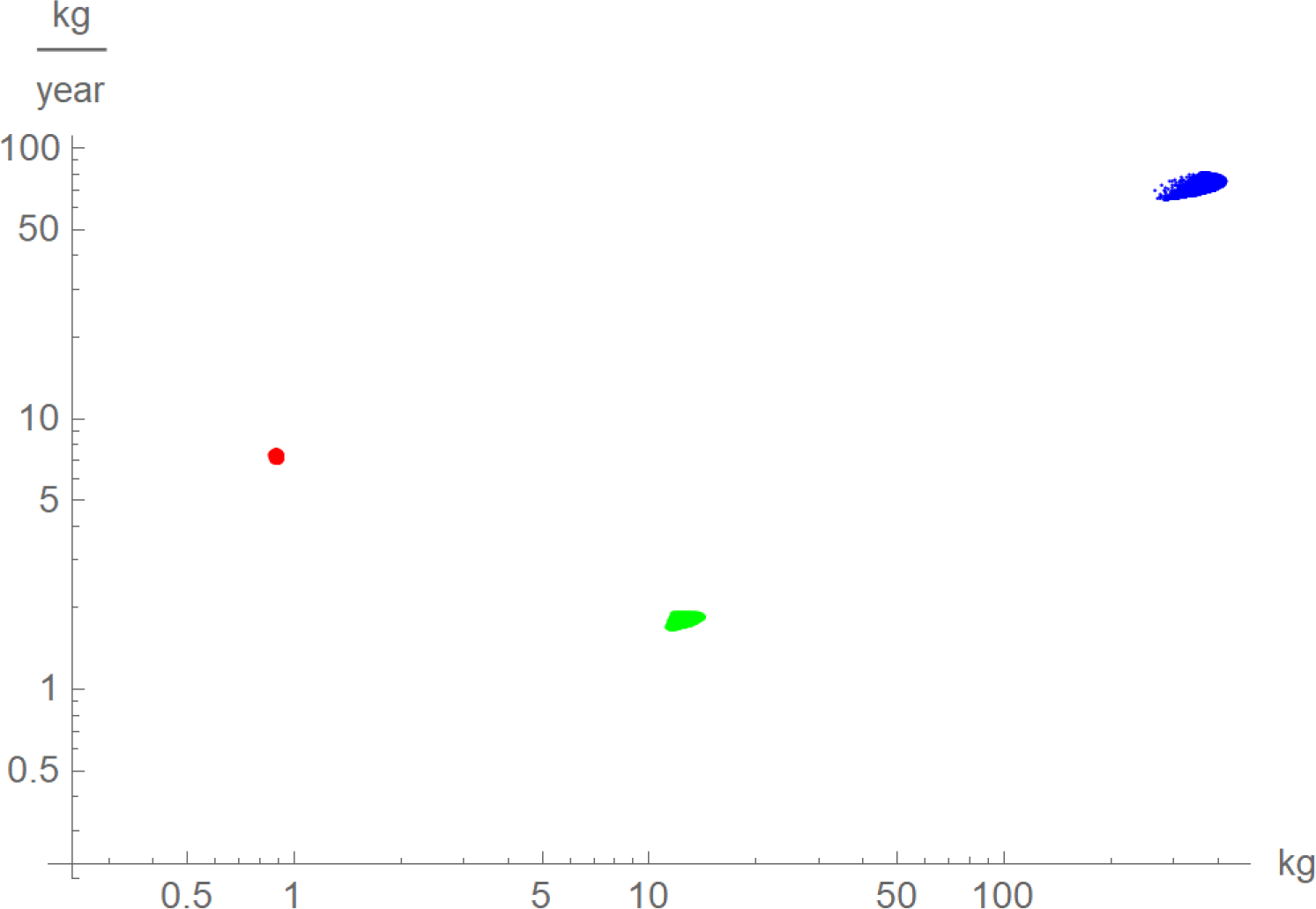
Log-log-plot of the maximal growth rate, *m’*, and mass at the inflection point for near-optimal growth curves *m*(*t*) for chicken (red), alligators (green) and dinosaurs (blue).

## 4. DISCUSSION

A large region of near-optimal exponents indicates that data may not carry enough information to differentiate between growth models. For the data about three species of dinosaurs from [16] only *Tenontosaurus* provided feasible data, while those for other species resulted in unreasonably large regions of near-optimal exponent-pairs (i.e. almost every growth model would be near-optimal). The paper therefore did not use these data. However, in view of the inherent uncertainties of estimating the mass of dinosaurs [5], it was surprising that one in three data-sets was suitable.

The definition of “full age” to define dimensionless coordinates was somewhat arbitrary. For, using 90% of the asymptotic mass was a compromise of avoiding excessive extrapolation (for some data the maximal observed mass was below the asymptotic mass) and the intent to cover most of the growth phase. Further, for different species the fraction *t*/*t*_*full*_ may correspond to different stages of their biological development. However, using this linear transformation was a convenient tool to combine data and growth curves of several species into one plot (Figures 2, 4 and 5). With respect to Figure 4, the faster growth of broiler chicken will also be observed for any nonlinear transformation of time that aims at a proper representation of biological development.

In Figure 3, the regions of near-optimal exponents displayed fuzzy boundaries and points close to the diagonal were not connected to the regions. This was caused by the optimization strategy, a high number of annealing steps for points next to the diagonal and few steps thereafter. (This speeded up computations.) However, despite these deficiencies the visualization of the near-optimal exponents verified the optimal character of the optimal exponent-pairs. As is evident from this figure, the optimal exponent-pairs were quite remote from the exponent-pairs for the named models which are more common in growth studies. However, in fish-biology it has long been accepted that exponent-pairs (*a*, *b*) with *a*< 1 and *b* = 1 might be better compatible with biological constraints for growth; e.g. the growth of gill surface area relative to mass growth [13]. Recently, also exponents *b* < 1 were considered as biologically meaningful [20]. Thus, the use of general exponent-pairs was also motivated by biological considerations.

## 5. CONCLUSION

While it is generally acknowledged that mass-at-age estimates for dinosaurs are highly uncertain, a data-set for *Tenontosaurus* allowed for the identification of a best fitting growth model within the comprehensive class of Bertalanffy-Pütter models (1). However, data uncertainty did not allow to conclude that the dinosaur-data would need a different exponent-pair (model) than modern alligators or birds. On the contrary, displaying the data in dimensionless coordinates did not indicate notable differences. Also, the best-fitting growth curves did barely differ. Yet, there was a difference in the relative growth rate, i.e. growth rate over mass. Thereby, modern broiler chicken grew much faster than dinosaurs or alligators and (keeping in mind the uncertainty of mass estimation) dinosaurs may grow faster than alligators. However, the growth rate is a measure that cannot be observed directly from the data; it is derived from a growth model and depends on what model is selected. This was demonstrated for the maximal growth rate, which varied considerably even for growth curves that fitted well to the data.

## REFERENCES

[1] Kahm, M., G. Hasenbrink, H. Lichtenberg-Frate, J. Ludwig, and M. Kschischo. 2010. Fitting Biological Growth Curves with R. Journal of Statistical Software, 33: 1–21.

[2] Aggrey, S.E. 2002. Comparison of Three Nonlinear and Spline Regression Models for Describing Chicken Growth Curves. Poultry Sciences, 81:1782–1788.

[3] Renner-Martin, K., N. Brunner, M. Kühleitner, W.G. Nowak, and K. Scheicher. 2018. Optimal and near-optimal exponent-pairs for the Bertalanffy-Pütter growth model. PeerJ 6: e5973.

[4] Lee, A.H., A.K. Huttenlocker, K. Padian, and H.N. Woodward. 2013. Analysis of Growth Rates. In Padian, K. and E.T. Lamm (eds.) Bone histology of Fossil Tetrapods: Advancing Methods, Analysis and Interpretation. UCLA Press: Berkeley, USA, pp. 217–251.

[5] Erickson, G.M. (2014). On Dinosaur Growth. Annual Review of Earth and Planetary Sciences, 42: 675–697.

[6] Brody, S. (1945) Bioenergetics and growth. Hafner Publ. Comp.: New York, NY, USA.

[7] Bertalanffy, L.v. (1949) Problems of organic growth. Nature 163: 156–158.

[8] Gompertz, B. (1832) On the Nature of the Function Expressive of the Law of Human Mortality, and on a New Mode of Determining the Value of Life Contingencies. Philos. Trans. R. Soc. London 123: 513–585.

[9] Richards, F.J. (1959) A Flexible Growth Function for Empirical Use. Journal of Experimental Botany 10: 290–300.

[10] Tjørve, M.C., and E. Tjørve. 2017. The use of Gompertz models in growth analyses, and new Gompertz-model approach: An addition to the Unified Richards family. PLoS ONE 12(6):e0178691.

[11] West, G.B., Brown, J.H., Enquist, B.J. (2001) A general model for ontogenetic growth. Nature 413: 628–631.

[12] Verhulst, P.F. (1838) Notice sur la loi que la population suit dans son accroissement, CorrespondenceMathematique et Physique (Ghent) 10: 113–121.

[13] Pauly D. (1981) The relationship between gill surface area and growth performance in fish: a generalization of von Bertalanffy’s theory of growth. Reports on Marine Research (Berichte der deutschen wissenschaftlichen Kommission für Meeresforschung) 28:25–282.

[14] Pütter, A. (1920) Studien über physiologische Ähnlichkeit. VI. Wachstumsähnlichkeiten. Pflügers Archiv für die Gesamte Physiologie des Menschen und der Tiere 180: 298–340.

[15] Bertalanffy, L.v. (1957) Quantitative laws in metabolism and growth. Quarterly Reviews of Biology 32: 217–231.

[16] Lee, A.H. and S. Werning. 2008. Sexual maturity in growing dinosaurs does not fit reptilian growth models. Proceedings of the National Academy of Sciences USA, 105: 582–587.

[17] Rootes, W.L., R.H. Chabreck, V.L. Wright, B.W. Brown, and T.J. Hess. 1991. Growth Rates of American Alligators in Estuarine and Palustrine Wetlands in Louisiana. Estuaries 14: 489–494.

[18] Renner-Martin, K., N. Brunner, M. Kühleitner, W.G. Nowak, and K. Scheicher. 2019. Best-fitting growth curves of the von Bertalanffy-Pütter type. Poultry Science, to appear, DOI: 10.3382/ps/pez122.

[19] Calder, W.A. III. 1985. Size, Function, and Life History. Harvard Univ. Press: Cambridge, USA.

[20] Pauly, D., and W.W.L. Cheung. 2017. Sound physiological knowledge and principles in modeling shrinking of fishes under climate change. Global Change Biology. Published online: DOI 10.1111/gcb.13831.

